# Relaxed targeting rules allow PIWI-clade Argonaute proteins to silence ever-mutating transposons

**DOI:** 10.1101/2022.08.04.502788

**Authors:** Ildar Gainetdinov, Katharine Cecchini, Joel Vega-Badillo, Ayca Bagci, Cansu Colpan, Amena Arif, Pei-Hsuan Wu, Phillip D. Zamore

**Affiliations:** RNA Therapeutics Institute and Howard Hughes Medical Institute, University of Massachusetts Chan Medical School; Worcester, MA, USA

## Abstract

In animals, piRNAs direct PIWI-clade Argonaute proteins to slice complementary transposon transcripts. Transposons can evade silencing through target site mutations. We report that PIWIs efficiently cleave transcripts only partially paired to their piRNA guide. Measurements of mouse PIWI protein affinity and cleavage rates for thousands of RNAs in vitro and in vivo show that PIWI slicing tolerates mismatches to any target nucleotide, including those flanking the scissile phosphate. Although piRNA 5’ terminal nucleotides accelerate target finding, they are dispensable for binding or catalysis—unlike AGO-clade Argonautes, which require uninterrupted siRNA:target pairing from the seed to the nucleotides past the scissile bond. PIWIs are thus better equipped than AGOs to target newly acquired or rapidly diverging endogenous transposons without recourse to novel small RNA guides.

## Introduction

In prokaryotes and eukaryotes, small RNA or DNA guides direct Argonaute proteins to fight viruses, plasmids, and transposons (*1–6*); regulate gene expression (*7–13*); and aid DNA replication (*14*). Animals produce two distinct types of Argonautes, AGOs and PIWIs. AGO clade proteins use small interfering RNAs (siRNAs, ~21 nt) or microRNAs (miRNAs, ~22 nt) to repress extensively or partially complementary transcripts (*7, 15*). AGOs initially find their targets via complementarity to a short 5’ region of their guide, the seed (nucleotides g2–g8; Fig. 1A); for miRNA-guided AGOs, seed complementarity suffices to repress the target RNA (*7, 16*). In contrast, PIWI-clade Argonaute proteins use PIWI-interacting RNAs (piRNAs, ~18–35-nt) as guides (*17*). While most eukaryotic genomes encode one or more AGO proteins, only animals make PIWI proteins (*18, 19*). With few exceptions, all animals use piRNAs to repress transposons (*17*). The ancestral mechanism of piRNA-guided transposon silencing is PIWI-catalyzed endonucleolytic cleavage (“slicing”) of complementary transposon RNAs in the cytoplasm (*20–23*). Moreover, piRNA production itself requires piRNA-directed slicing of piRNA precursor transcripts (*20, 21, 24–26*). In some animals, piRNAs also direct nuclear PIWI proteins to nascent transposon transcripts to silence transcription via repressive histone marks or DNA methylation (*27–31*).

**Fig. 1.**
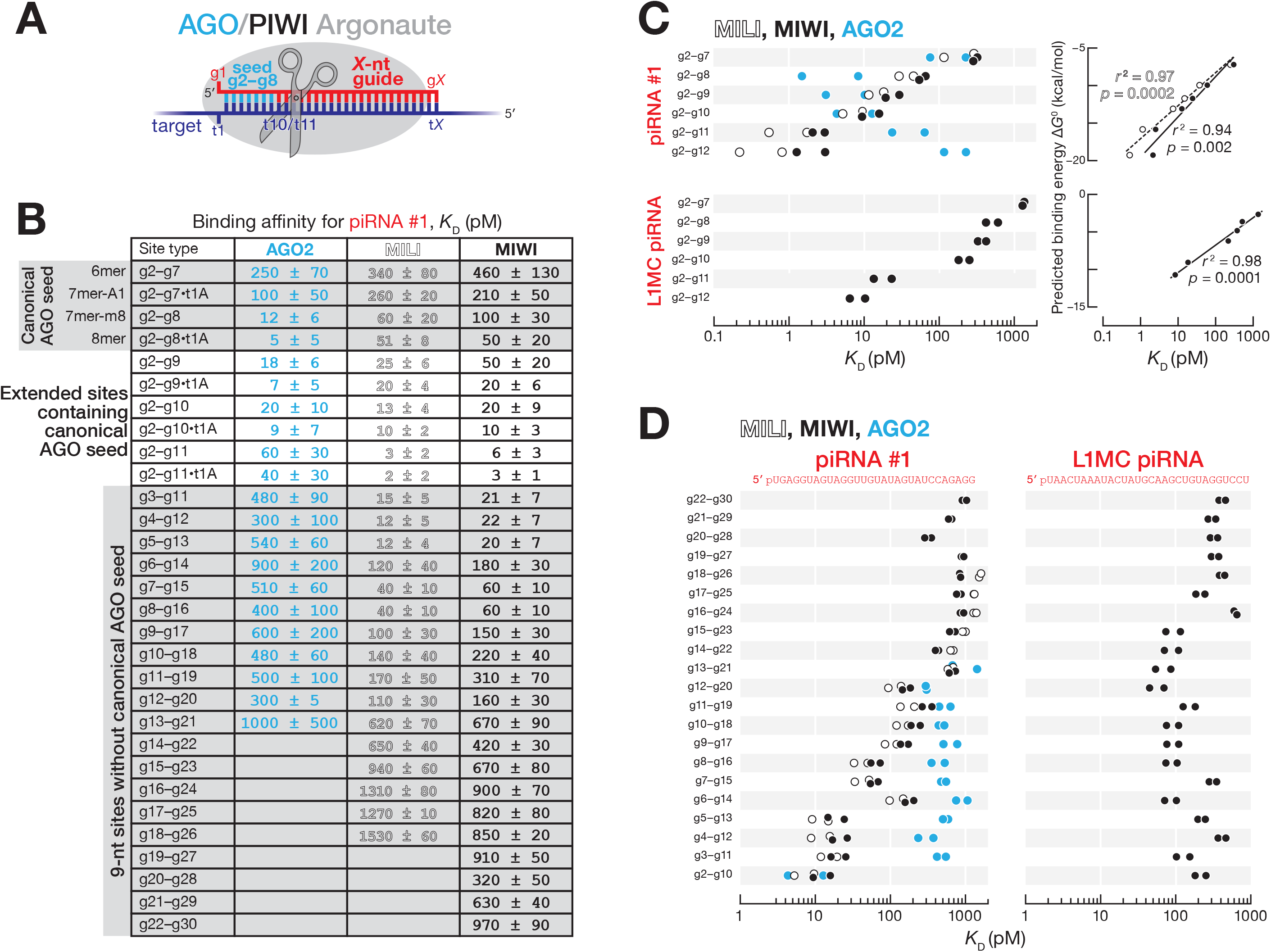
Mouse PIWI proteins bind sites with and without canonical seed pairing. (**A**) Small RNA guides direct eukaryotic Argonaute proteins to complementary targets. (**B**) Binding affinities (*K*_D_) of AGO2, MIWI, and MILI loaded with piRNA #1 for canonical and non-canonical target sites. Mean and the range of the data from two independent trials are shown. (**C**) Left, AGO2, MIWI, and MILI binding affinities (*K*_D_) for targets contiguously paired from nucleotide g2. Data are from two independent trials. Right, relationship between *K*_D_; and predicted binding energy Δ*G*^0^. *K*_D_; is the mean of two independent trials. Goodness-of-fit for linear regression (*r*^2^) and *p* value for Spearman’s correlation are shown. (**D**) AGO2, MIWI, and MILI binding affinities for 9-nt complementary stretches contiguously paired from all guide nucleotides. Data are from two independent trials. AGO2 data are from Ref (*39*).

siRNAs direct AGO proteins (*32*) and piRNAs direct PIWI proteins (*22*) to hydrolyze the phosphodiester bond joining the target nucleotides opposite guide nucleotides g10 and g11 (i.e., t10 and t11; Fig. 1A). Unlike siRNA-directed, AGO-catalyzed transcript cleavage, target slicing by PIWI proteins requires the auxiliary factor GTSF1/Asterix (*33*). GTSF1 accelerates ~10–100-fold the otherwise slow target cleavage by PIWIs, likely by stabilizing the catalytically competent conformation of PIWI proteins (*33*). Why PIWI slicing evolved to require an auxiliary protein is unknown.

Here, we report that the requirements for guide:target complementarity are relaxed for the mouse PIWI proteins MILI (PIWIL2) and MIWI (PIWIL1) compared to AGO proteins. PIWIs bind RNAs both with and without complementarity to the canonical 5’ seed of their guide. Both in vitro and in vivo, PIWI-catalyzed slicing requires at least 16 contiguously paired nucleotides; longer extents of complementarity tolerate guide:target mismatches at essentially any position. Unlike AGO proteins, guide pairing to any target nucleotide, including those that flank the scissile phosphate (t10 and t11), is dispensable for efficient slicing. Although pairing to ≥ 4 piRNA 5’ terminal nucleotides facilitates target finding, in vitro and in vivo abundant piRNAs direct slicing of targets lacking 5’ complementarity. Notably, the minimum 16-nt stretch of complementarity that licenses piRNA-guided target cleavage suffices to distinguish host transcripts from transposon RNAs. These findings suggest that the catalytic properties of PIWI proteins evolved to prevent transposons from escaping piRNA silencing via mutation while simultaneously retaining sufficient specificity to spare “self” transcripts from inappropriate repression.

### PIWI proteins bind efficiently to sites both with and without canonical seed pairing

piRNAs have been proposed to direct MIWI to bind and regulate mRNA expression via the same mechanism by which miRNAs guide AGOs to its targets (*34, 35*). In this miRNA-like binding mode, base pairing to the canonical AGO seed sequence (guide nucleotides g2–g8; Fig. 1A) mediates the search for complementary sites and suffices to tether the Argonaute protein to its target RNA. We used RNA Bind-’n-Seq (RBNS) (*36–39*) to measure the affinity of piRNA-guided PIWI Argonautes for a library of 20-nt-long random sequences (fig. S1A). We incubated the target RNA library with either purified MILI or MIWI loaded with a 5’ monophosphorylated synthetic RNA (26 nt for MILI, 30 nt for MIWI; fig. S1B) (*33*) and isolated and sequenced RNAs bound to the piRNA•PIWI-protein complex (piRISC). The sequencing data was analyzed using an approach that estimates the affinity (*K*_D_) of piRISC for each binding-site type (*39*).

Binding of MILI or MIWI to RNAs with canonical AGO seed sites was weaker than for mouse AGO2 (Fig. 1B). Compared to AGO2, MILI and MIWI affinity was ~10-fold lower for an 8mer and ~5-fold lower for a 7mer-m8, the two canonical site types most effectively repressed by miRNA-guided AGO proteins (*7*): for piRNA #1, 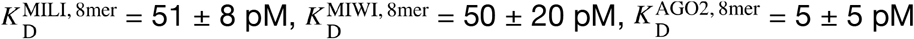 (Fig. 1B). The intracellular concentration of the most abundant piRNAs (~19 nM) is less than the most abundant miRNAs in mouse primary spermatocytes (~24 nM) (*40*), suggesting that the weaker affinity of PIWI proteins for the canonical seed sites will result in lower occupancy of such targets. Our data therefore disfavor a model in which PIWI proteins find and productively regulate targets via 7–8 nt canonical seed pairing.

For AGO2, extending pairing beyond the canonical seed does not increase the affinity of RISC for a target. In contrast, extending guide:target complementarity from g2–g8 to g2–g10 increased piRISC affinity for targets ~five-fold: 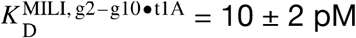 vs 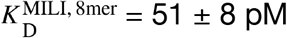, and 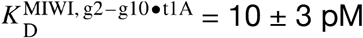 vs 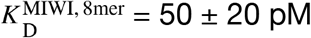 compared with 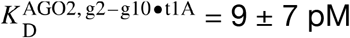 vs 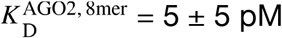 (Figs. 1B and 1C). For both a piRNA of synthetic sequence (piRNA #1) and a piRNA found in vivo (L1MC piRNA), the binding affinity of MILI and MIWI piRISC increased linearly with increasing predicted base pairing energy, Δ*G*° (Fig. 1C). Thus, MILI and MIWI, like EfPiwi (*41*), require a longer extent of guide:target base pairing to approach the affinity of AGO2 RISC for a canonical seed match.

Unlike AGOs, MILI and MIWI bound tightly to sites both with and without full pairing to the canonical seed (Figs. 1B and 1D, and fig. S1C). For instance, MILI and MIWI bound to targets with complementarity to the guide starting at g2 or g5 with indistinguishable affinities: 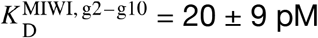 vs 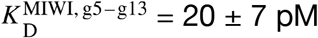, and 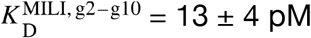 vs 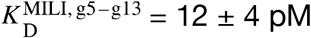 (Figs. 1B and 1D). In contrast, AGO2 bound sites lacking seed pairing 10–100-fold more weakly than to those with a seed match (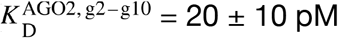 vs 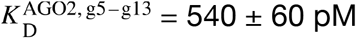; Figs. 1B and 1D).

Curiously, MIWI directed by a piRNA antisense to the mouse L1MC retrotransposon bound more tightly to a 9-nt site with no seed match 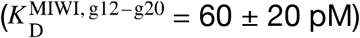 than to 9-nt sites with partial seed complementarity (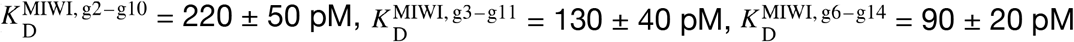; Fig. 1D). We conclude that MILI and MIWI are more flexible than AGO2 in the types of sites they can bind, but require longer complementarity to achieve high target affinity.

### Pairing to piRNA 3’ sequence permits slicing of partially complementary targets

The modal length of piRNAs is 5–10 nt longer than that of siRNAs (26–31 nt vs ~21 nt) (*4, 42–45*), yet PIWI proteins do not require pairing to these additional 3’ nucleotides to cleave a target RNA (*22, 33, 41*). In vitro, 16–21-nt long contiguous complementarity sufficed for MILI and MIWI to reach their maximum endonuclease rate (fig. S2A). Moreover, MILI and MIWI, directed by 21-nt guides, cleave targets as efficiently as when loaded with full-length piRNAs (*33*). Nonetheless, we find that extending pairing beyond piRNA nucleotide g20 allows PIWI proteins to tolerate guide:target mismatches.

We used a high-throughput approach, Cleave-’n-Seq (CNS) (*46*), to determine the rates of cleavage for thousands of target variants (fig. S2B). We incubated purified MILI or MIWI piRISC (1 nM) and the PIWI auxiliary factor GTSF1 (500 nM) with a library containing ~7,700–10,400 ~30-nt long target RNAs for different lengths of time (60 sec to 16 h). Uncleaved RNAs were reverse-transcribed, sequenced, and their abundance at each time point used to determine their pre-steady-state cleavage rate, *k* (fig. S2B). We conducted CNS using four different piRNA guides for both MILI and MIWI (fig. S1B).

Consistent with the idea that additional complementarity to piRNA 3’ nucleotides accelerates cleavage of imperfectly paired targets, a mismatch between g2 and g20 decreased the median *k* ~3.3-fold when all nucleotides after g20 were unpaired, but only ~1.4-fold when g21—g25 were also base paired (Fig. 2A). Two mismatches between g2 and g20 caused a ~17-fold median reduction in *k* for targets with no pairing beyond g20, but just ~3-fold when g21–g25 were paired (Fig. 2A). Thus, endonucleolytic cleavage by MILI or MIWI does not require target pairing to piRNA 3’ sequences, but such extended complementarity readily compensates for guide:target mismatches.

**Fig. 2.**
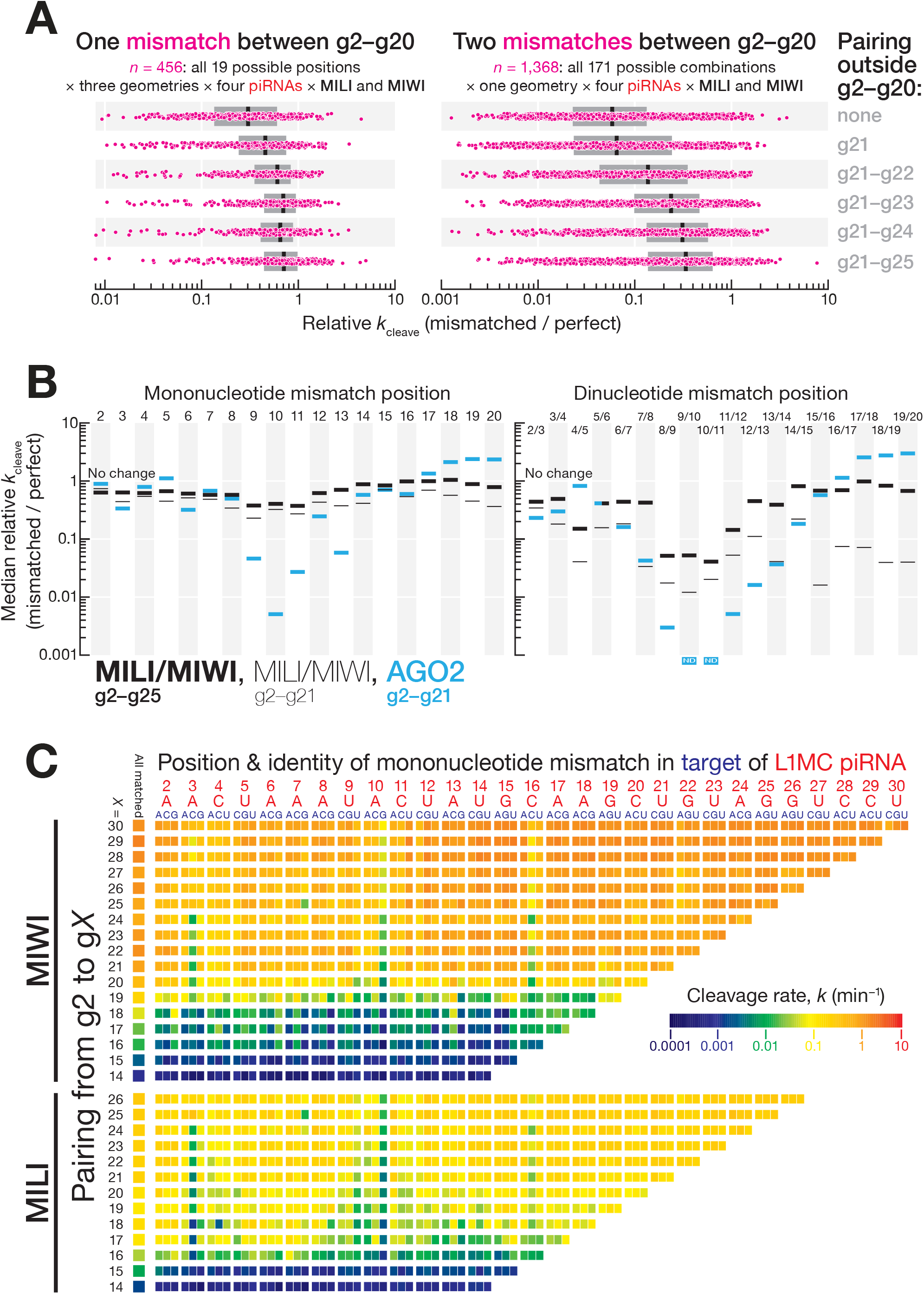
Mouse PIWI slicing tolerates mismatches with any target nucleotide. (**A**) Change in pre-steady-state cleavage rate for one or two mismatches between g2–g20. Box plots show inter-quartile range (IQR) and median. (**B**) Median change in presteady-state cleavage rate for one or two consecutive mismatches between g2–g20 for contiguous g2–g21 or g2–g25 pairing. All data are in S3A. (**C**) MILI and MIWI pre-steady-state cleavage rates for targets of L1MC piRNA containing a single unpaired nucleotide.

Argonautes secure target nucleotide t1 in a pocket that often displays specific nucleotide preferences. Although MILI and MIWI showed a slight binding preference for t1A (≤ 3-fold stronger affinity compared to t1B; Fig. 1B and fig. S2C), target cleavage showed no t1 preference (fig. S2A).

### PIWI slicing tolerates mismatches to any target nucleotide including those flanking the scissile bond

AGO2-catalyzed cleavage requires uninterrupted pairing to siRNA nucleotides g9–g13 (*46*). In contrast, MILI and MIWI slicing tolerated mismatches at any position within the region of complementarity (Figs. 2B and 2C, figs. S3, S4, and S5).

For g2–g21-paired targets of AGO2, a single mononucleotide mismatch at g9, g10, g11, or g13 decreased median *k* 7–200-fold (Fig. 2B and fig. S3A) (*46*). For the same extent of pairing, the median reduction in *k* for MILI and MIWI piRISC was < 4-fold for a mononucleotide mismatch at any position between g2 and g20 (Fig. 2B and fig. S3A). Consistent with the compensatory effect of pairing to piRNA 3’ nucleotides (Fig. 2A), guide:target complementarity from g2 to g25 reduced the median impact of a mononucleotide mismatch to ≤ 2.7-fold at any position between g2 and g20 (Fig. 2B and fig. S3A). Among mismatch types, GU wobbles had the smallest impact on slicing rates, decreasing *k* just ~1.2-fold (fig. S3B).

Unlike AGO2 RISC, pairing to target nucleotides adjacent to the scissile phosphate was dispensable for target slicing by piRISC (Figs. 2B and 2C, figs. S3, S4, and S5). Loss of pairing to either t10 or t11 in a g2–g21 match decreased the cleavage rate by ≥ 10-fold for only 11 of 48 tested targets for four different piRNAs bound to MILI or MIWI, and the median decrease was 2.5-fold for unpaired t10 (inter-quartile range, IQR: 1.6–9.1) and 2.7-fold for unpaired t11 (IQR: 1.7–5.2) (Fig. 2C). Purine-purine mismatches at these positions appeared to be least well tolerated by piRISC, perhaps because their greater bulk is less well accommodated within the PIWI catalytic center (fig. S3C). MILI- and MIWI-catalyzed slicing was detectable even when both t10 and t11 were unpaired: the median decrease in *k* was 25- and 50-fold for g2–g25 and g2– g21 pairing, respectively (Fig. 2B and fig. S3A). In contrast, no target cleavage by AGO2 RISC was detected for the same dinucleotide mismatch at t10–t11 (Fig. 2B and fig. S3A). We conclude that MILI and MIWI, unlike AGO2, can efficiently cleave partially paired RNAs with mismatches anywhere in a target site.

### piRNAs direct cleavage of partially complementary targets in vivo

In mice, piRNAs guide MILI and MIWI to slice complementary transposon transcripts, mRNAs, or long non-coding transcripts (lncRNAs) (*8, 9, 11, 12, 22, 23*). As we observed for purified piRISC, in vivo in mouse primary spermatocytes, piRNAs direct MILI and MIWI to cleave targets with as few as 15–19-nt complementary nucleotides.

Mouse primary spermatocytes produce a class of MILI- and MIWI-loaded piRNAs called pachytene piRNAs, which first appear at the pachytene stage of meiosis (*42, 43, 45, 47*). Because endonucleolytic cleavage by Argonaute proteins leaves a 5’-monophosphate (*48*), we sequenced 5’-monophosphorylated RNAs from FACS-purified mouse primary spermatocytes to identify potential 3’ cleavage products generated in vivo by MILI and MIWI (Fig. 3A). Restricting our analysis to pachytene piRNAs, > 80% of which derive from non-repetitive sequences, ensured unambiguous assignment of piRNAs to candidate cleavage products. To identify those RNAs corresponding to 3’ cleavage products generated by piRNA-directed slicing, we searched for 5’ monophosphate-bearing RNAs present in control C57BL/6 mice, but whose abundance was reduced ≥ 8-fold in a triple mouse mutant lacking all piRNAs from three major pachytene piRNA-producing loci on chromosomes 2, 9, and 17: 2-qE1-35981[+]; 9-qC-31469(-),10667(+); 17-qA3.3-27363(-),26735(+) (*11, 47*). For simplicity we refer to these loci as *pi2, pi9*, and *pi17*. The *pi2^−/−^; pi9^−/−^; pi17^−/−^* triple mutation removes ~22% of all pachytene piRNAs (Fig. 3A and fig. S6).

**Fig. 3.**
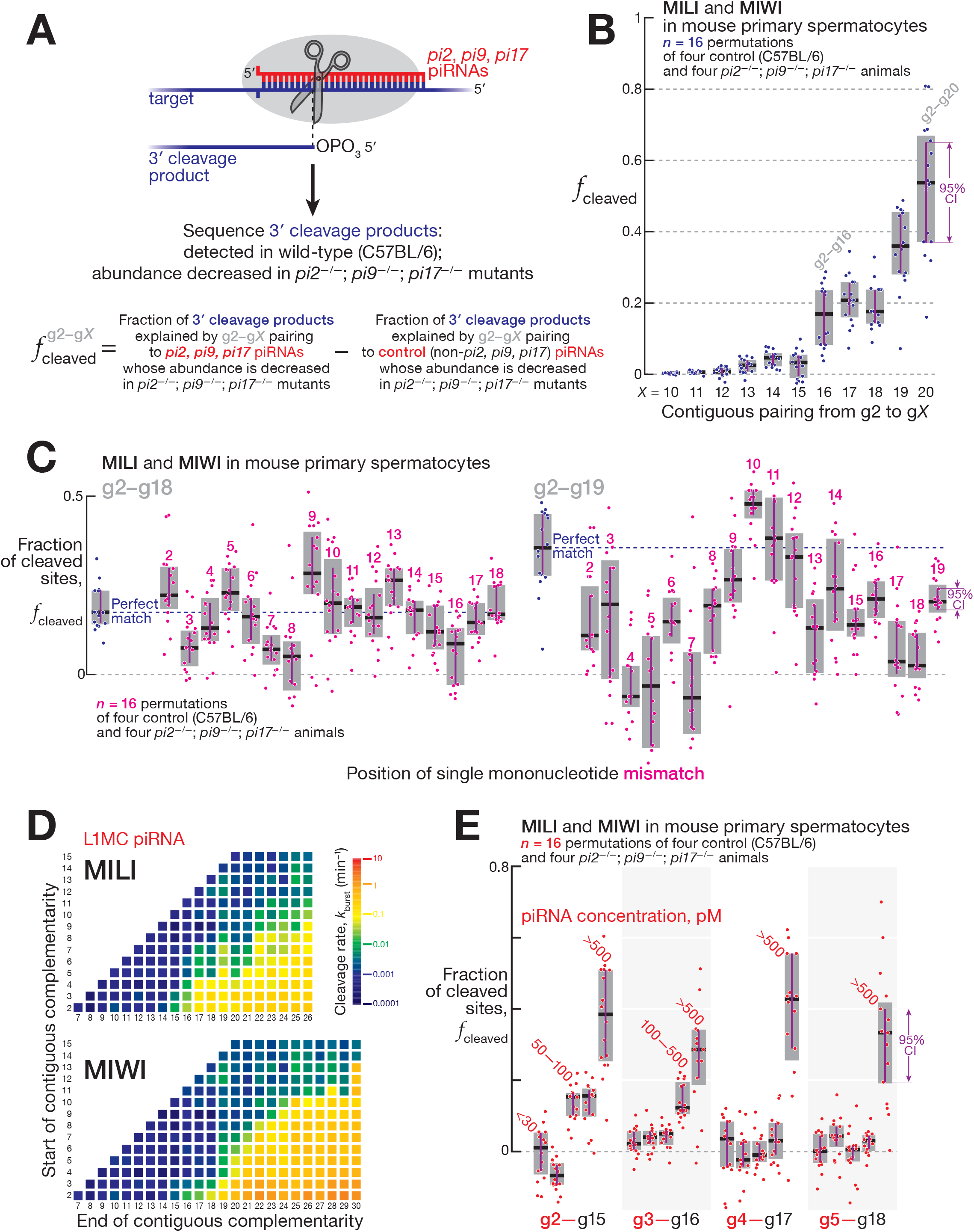
Mouse PIWI proteins cleave partially complementary targets in vivo. (**A**) Strategy to identify 3’ cleavage products of piRNA-guided PIWI-catalyzed slicing and measure the fraction of targets cleaved by PIWI proteins in FACS-purified mouse primary spermatocytes. (**B**) Fraction of cleaved targets in FACS-purified mouse primary spermatocytes for contiguous pairing from nucleotide g2. Box plots show IQR and median; 95% confidence interval was calculated with 10,000 bootstrapping iterations. (**C**) Fraction of cleaved targets in FACS-purified mouse primary spermatocytes for pairing containing a single-nucleotide mismatch. Box plots show IQR and median; 95% confidence interval was calculated with 10,000 bootstrapping iterations. (**D**) MILI and MIWI pre-steady-state cleavage rates in vitro for all possible stretches of ≥ 6-nt contiguous pairing starting from nucleotides g2–g15 of L1MC piRNA. (**E**) Fraction of cleaved targets in FACS-purified mouse primary spermatocytes for 14-nt contiguous pairing starting from nucleotides g2–g5. Data are binned by piRNA intracellular concentration. Box plots show IQR and median; 95%confidence interval was calculated with 10,000 bootstrapping iterations.

Among the 5’-monophosphorylated RNAs detected in the control C57BL/6 mice, we selected candidate cleavage products whose production could be explained by a *pi2, pi9*, or *pi17* piRNA directing cleavage between nucleotides t10 and t11. For each pattern of piRNA:target complementarity, e.g., g2–g*X* pairing with t2–t*X*, we calculated the fraction of cleavage product candidates whose abundance was reduced ≥ 8-fold in *pi2^−/−^; pi9^−/−^; pi17^−/−^* triple mutant primary spermatocytes. We denote this fraction as 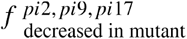. A pairing configuration that can support target slicing is predicted to have a high fraction of such candidate 3’ cleavage products present in the control but reduced or absent in the triple mutant mice (Fig. 3A).

To account for sampling error arising from the short-lived nature of 5’-monophosphorylated fragments in vivo, we identified all 5’-monophosphorylated RNAs in C57BL/6 explained by piRNAs *not removed* in *pi2^−/−^; pi9^−/−^; pi17^−/−^* mice (control piRNAs), and then calculated the fraction of these RNAs reduced by ≥ 8-fold in *pi2^−/−^*; *pi9^−/−^; pi17^−/−^* animals. The fraction of cleaved targets for each pairing arrangement g2–g*X* was then calculated as the observed signal minus the sampling error (Fig. 3A).

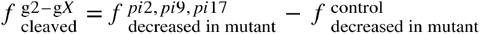

Cleavage by MILI or MIWI is indistinguishable in our data, thus 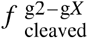 corresponds to the sum of targets sliced in mouse primary spermatocytes by both PIWI proteins.

These analyses showed that PIWI-catalyzed cleavage was detected in primary spermatocytes for piRNA:target base pairing as short as g2–g16: median 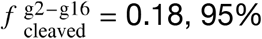 confidence interval (CI): [0.08, 0.23] (Fig. 3B). The efficiency of piRNA-directed target cleavage increased with longer complementarity: e.g., median 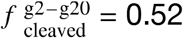 (CI: [0.37, 0.65], Fig. 3B). (Pairing longer than g2–g20 contained too few cleaved data points to measure the corresponding 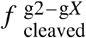.)

When piRNA:target RNA complementarity was ≥ 17-nt long, single mismatches were tolerated at most positions (Fig. 3C and fig. S7A). For example, for g2–g18 complementarity, median *f*_cleaved_ or perfect pairing (0.18, CI: [0.14, 0.23]) was similar to g2–g18 matches bearing a single nucleotide mismatch at positions g2, g5, g6, g9–g14, or g17–g18 (0.15–0.25; Fig. 3C). For targets complementary to piRNA nucleotides g2– g18 or g2–g19, pairing to t10 and t11, the target nucleotides flanking the scissile phosphate, was dispensable for slicing (Fig. 3C).

These in vivo data mirror the results of our biochemical assays showing that the median decrease in cleavage rate was ≤ 4-fold for a mononucleotide mismatch at any position between g2 and g20 (Fig. 2B and fig. S3A). We conclude that a wide variety of piRNA:target pairing patterns can efficiently direct MILI and MIWI to cleave targets, unlike the relatively limited pairing configurations tolerated by AGO-clade Argonautes.

### Highly abundant piRNAs guide slicing of targets lacking complementarity to the canonical 5’ seed

AGO2-catalyzed slicing of sites contiguously paired from g4 or g5 (i.e., without canonical seed pairing) has been observed in vitro but not detected in vivo (*46, 49*). By contrast, our data identify piRNA-directed cleavage in vivo in mouse primary spermatocytes of targets whose pairing to the guide starts at g3, g4, or g5.

In vitro, MILI and MIWI did not require target pairing to piRNA 5’ terminal nucleotides for binding (Figs. 1B and 1D) or slicing (Fig. 3D and fig. S7B). Similarly, we detected in vivo piRNA-directed cleavage of targets lacking complementarity to nucleotides g2–g4 (Fig. 3E). Pre-organization of seed nucleotides g2–g7 in an A-form-like helix accelerates target finding by AGO proteins (for the let-7a 8mer, 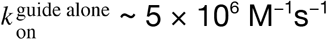 vs 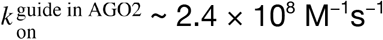) (*50–52*). PIWI proteins preorganize only nucleotides g2–g4 (*41, 53*), suggesting slower target finding when piRNA:target complementarity begins after nucleotide g4. In theory, high piRISC concentration could compensate for a slower target-finding rate constant (*k*_on_). In vivo, piRNA concentrations vary widely, and ~1,500 piRNAs are present in mouse primary spermatocytes at ≥ 500 pM (Figure S7C). We observed piRNA-directed cleavage of targets with contiguous 14-nt pairing beginning at g4 (median 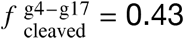, CI: [0.29–0.56]) or g5 (median 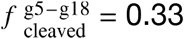, CI: [0.2–0.4]) only for highly abundant piRNAs (≥ 500 pM). Cleavage of targets paired from g3 was detectable for piRNAs whose in vivo concentration was ≥ 100 pM, and targets paired from g2 were sliced by piRNAs present at ≥ 50 pM (Fig. 3E).

We conclude that, because pairing to piRNA 5’ terminal nucleotides is dispensable for both target finding and slicing, piRISC efficiently cleaves targets lacking full complementarity to the canonical 5’ seed.

### piRNA-directed slicing does not tolerate mononucleotide target insertions or deletions

As observed for AGO2 (*46*), mononucleotide target insertions between t9–t15 slowed MILI- and MIWI-catalyzed cleavage in vitro by ≥ 10-fold (fig. S8A). For both AGO2 (*46, 54*) and the Piwi protein from the sponge *Ephydatia fluviatilis* (*Ef*Piwi) (*41*), target nucleotides t9–t15 face the protein surface making insertions likely to distort the catalytic center.

Single-nucleotide target deletions between t6–t15 were also poorly tolerated (≥ 10-fold lower *k* in vitro for MILI and MIWI, relative to a fully complementary target; fig. S8B). Such target sequence deletions result in mononucleotide bulges in the piRNA guide. Like mammalian AGO2 (*54*), *EfPiwi* restricts piRNA nucleotides g6–g10 to the protein’s central cleft (*41*), and a mononucleotide piRNA bulge between t6–t10 is unlikely to fit in this narrow furrow, potentially explaining why MILI and MIWI do not tolerate such target deletions. In contrast, single nucleotide deletions between t11–t15 create solvent-facing, mononucleotide loops of piRNA nucleotides g11–g15 that are predicted to be accommodated. Consistent with this idea, AGO2 tolerates target deletions between t11–t15 (*46*), yet MILI and MIWI did not (fig. S8B). Perhaps piRNA guide bulges between t11–t15 disrupt interactions with GTSF1.

In agreement with the in vitro data, PIWI-catalyzed cleavage in mouse primary spermatocytes was impaired in the presence of target insertions or deletions in the center of the piRNA:target duplex (fig. S8C).

### piRNA concentration and target affinity predict the efficiency of piRNA-guided cleavage in vivo

To identify the factors predicting effective piRNA slicing in vivo, we used the logistic regression classifier approach (*55, 56*). In vivo cleavage data were fit to a logistic function representing the probability of piRNA-guided cleavage, *P*(cleaved), determined by 35 variables (*X*_1_, *X*_2_, … *X*_35_): the presence or absence of pairing with each guide nucleotide between g2–g25, total number of paired nucleotides, predicted binding energy, piRNA abundance, target site location in the transcript (5’UTR, ORF, 3’UTR, or lncRNA), and the identity of target nucleotide t1 (Fig. 4A). The coefficient for each variable in the fitted logistic decision function (*β*_1_, *β*_2_, … *β*_35_) estimates the importance of each feature.

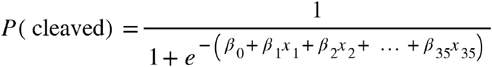

**Fig. 4.**
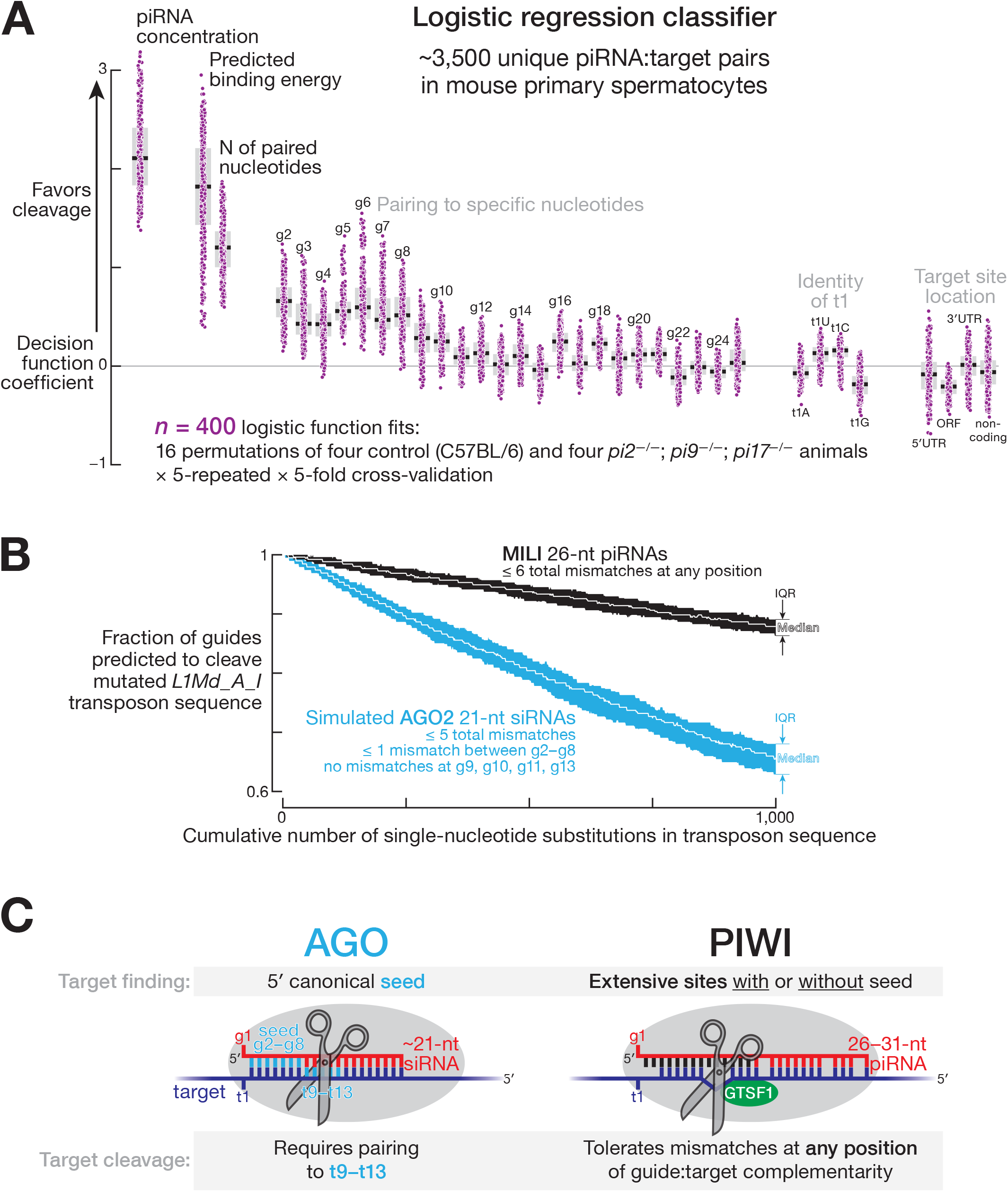
Determinants of PIWI slicing in vivo. (**A**) Decision function coefficients for 400 logistic function fits (regression models) using ~3,500 unique piRNA:target pairs detected in mouse primary spermatocytes. Box plots show IQR and median. (**B**) Number of piRNAs and siRNAs predicted to cleave mutated versions of the L1Md_A transposon sequence. Data are median and IQR from 100 independent simulations. (**C**) AGO and PIWI proteins use different rules to find and slice targets.

We selected ~3,500 unique pairs of *pi2, pi9, p17* piRNAs and target sites for which 5’-monophosphorylated 3’ cleavage products were both detected in the control C57BL/6 mice and had ≥ 19 nucleotides paired between g2–g25. Target sites were considered cleaved if the abundance of the 3’ cleavage products decreased ≥ 8-fold in the triple *pi2^−/−^; pi9^−/−^; pi17^−/−^* mutant compared to the control mice; all other target sites were assigned as uncleaved. To test if the logistic regression models created with the *pi2, pi9, pi17* piRNA data could predict cleavage by *non-pi2, pi9, pi17* piRNAs, we generated independent datasets from mice with a mutation disrupting a pachytene piRNA-producing locus on chromosome 7, *pi7* (7-qD2-24830(-)11976(+); fig. S6): > 99% of piRNAs eliminated in the *pi7^−/−^* deletion are not found in *pi2, pi9*, or *pi17*. The performance of the logistic function fitted to *pi2^−/−^; pi9^−/−^; pi17^−/−^* data was similar when tested with either *pi2^−/−^; pi9^−/−^; pi17^−/−^* (fig. S9A) or *pi7^−/−^* mutant data (fig. S9A).

The most predictive features, i.e., highest median coefficients in the logistic decision function, were piRNA concentration (+2.11), predicted energy of piRNA:target base pairing (+1.82), and the total number of paired nucleotides (+1.20; Fig. 4A). These results show that, in vivo, piRISC behaves as a conventional enzyme: its concentration and substrate binding strength determine the efficacy of target cleavage. A high aggregate number of targeting piRNAs was also recently shown to be required for potent transcriptional silencing (*57*).

Consistent with the idea that PIWI slicing does not rely on complementarity to specific target nucleotides (Figs. 2B and 3C), the median decision function coefficients were ≤ +0.6 for guide:target pairing at all positions (Fig. 4A). Extensive complementarity anywhere in the piRNA 5’ half appears to initiate target binding (Figs. 1B and 1D), and highly abundant piRNAs (≥ 500 pM) even direct slicing of targets without pairing to positions g2–g4 (Fig. 3E). In agreement with these data, logistic function coefficients for matches to g2–g10 were higher than those for pairing to other nucleotides ([+0.25, +0.6] vs [-0.12, +0.25]; Fig. 4A).

The features with the lowest median coefficients were the identity of nucleotide t1 and location of the target site in a transcript (−0.18 to +0.13; Fig. 4A), suggesting that these factors are not rate-determining for piRNA-guided cleavage in vivo.

Together, these analyses show that simple biochemical principles suffice to predict efficient piRNA-directed cleavage in cells. First, piRNA concentration determines how frequently a target encounters piRISC and thus the concentration of piRISC•target complex. Second, tighter guide:target base pairing (binding energy) likely extends the lifetime of the piRISC•target complex, increasing the likelihood of cleavage.

### Implications of relaxed piRNA targeting rules for transposon silencing

Because PIWI-catalyzed slicing does not depend on pairing to a specific piRNA nucleotide position, target mutations are predicted to be better tolerated by PIWIs compared to AGO proteins. Our computational simulations estimated that, when a transposon sequence mutates but the small RNA repertoire does not change, the number of guides capable of directing target slicing decreases ~4-fold slower for PIWI than AGO proteins (Fig. 4B and fig. S9B). Using the established mouse germline mutation rates (*58*), we simulated 1000 rounds of single-nucleotide substitutions in the LINE1 consensus sequences, excluding non-synonymous mutations in ORFs. At each round we recorded the number of embryonic testicular piRNAs or siRNAs (simulated using piRNA 21-nt prefixes) expected to productively slice the mutated transposons. The number of MILI-loaded piRNAs capable of cleavage decreased at 0.01% of guides per single-nucleotide substitution in LINE1 elements, while the number of simulated AGO2-loaded siRNAs decreased at 0.04% of guides per mutation in transposon (Fig. 4B and fig. S9B).

## Discussion

Our data highlight several distinct features of MILI and MIWI that set them apart from the AGO clade of Argonaute proteins.

First, PIWI proteins do not limit target finding to the 7-nt, 5’ canonical seed. The central cleft that cradles the guide RNA is wider in PIWIs than in AGOs (*41*), perhaps allowing PIWI proteins to productively use piRNA nucleotides 3’ to g8 to initiate pairing with targets. At high concentrations piRISC efficiently binds and slices RNAs unpaired to nucleotides g2–g4, so targeting capacity is similar for piRNAs whose 5’ ends are several nucleotides apart along a piRNA precursor transcript. This observation explains why piRNAs can tolerate a surprisingly high degree of 5’ heterogeneity, an intrinsic feature of phased (“trailing”) piRNA biogenesis (*24–26, 40, 59*). In contrast, miRNA 5’-isoforms have distinct target repertoires (*7*), and any change in the 5’ position of an siRNA duplex can invert which strand becomes a guide for an AGO protein (*60, 61*).

Second, piRNA-directed slicing tolerates mismatches to any nucleotide of the piRNA guide. We propose that, because piRNAs are generally longer than siRNAs, PIWIs can extend the lifetime of the piRISC•target complex through compensatory pairing to piRNA 3’ sequences, allowing piRISC to tolerate multiple guide:target mismatches. This hypothesis predicts that cleavage by mammalian PIWIL3 proteins, which are guided by unusually short, ~19-nt piRNAs (*62, 63*), will require full complementarity between their guides and targets. (PIWIL3 is produced in the ovaries of many mammals, but not mice (*64–67*).)

Third, the ability of piRNAs to direct cleavage of imperfectly complementary RNAs only when the extent of complementarity is longer than 16 contiguous base pairs (fig. S2A) (*22, 33, 41*) may explain the rapid evolution of the piRNA target repertoire. Our analyses suggest that a 16-nt contiguous match should suffice to prevent inappropriate targeting of mRNAs and other “self” transcripts. We determined the fraction of all mouse mRNAs and lncRNAs that contain at least one *k*-mer from the consensus sequences of transpositionally active families of mouse LTR or LINE transposons (*68, 69*). For *k* ≥ 16, fewer than 3% of mRNAs and lncRNAs shared at least one *k*-mer with transposon sequence (fig. S10, top). We obtained similar results when these analyses were conducted for the intron sequences of the same mouse transcripts (fig. S10, bottom), suggesting that strong negative selection against transposon-derived ≥ 16-mers in mRNAs and lncRNAs is unlikely.

GTSF1 has been proposed to stabilize the catalytically competent geometry between the target and the PIWI catalytic center (*33*). For AGO proteins, perfect pairing between an siRNA and its target moves target nucleotides t10 and t11 into the endonuclease active site. We speculate that partial pairing between piRNA and target suffices for MILI- and MIWI-catalyzed target cleavage because GTSF1, rather than guide:target base pairing, positions the scissile phosphate in the PIWI catalytic center (Fig. 4C).

---

**Fig. S1.**
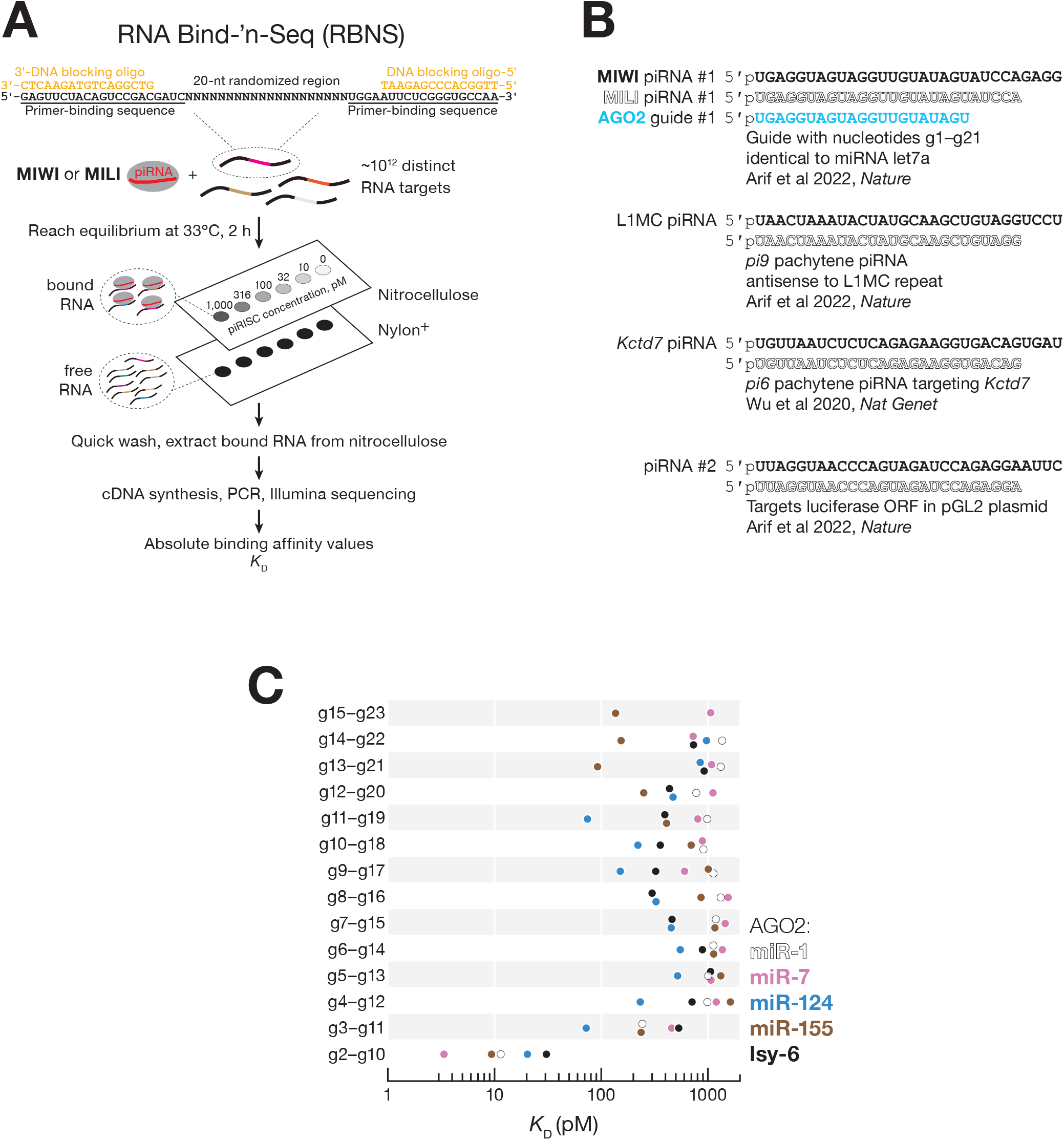
Binding Affinities of Argonautes, Related to Fig. 1. (A) Overview of RNA Bind-’n-Seq. (B) Sequences of small RNA guides used in this study. (C) AGO2 affinities for 9-nt complementary stretches contiguously paired from all guide nucleotides for miR-1, miR-7, miR-124, miR-155, and lsy-6. Data are from Ref (*38*).

**Fig. S2.**
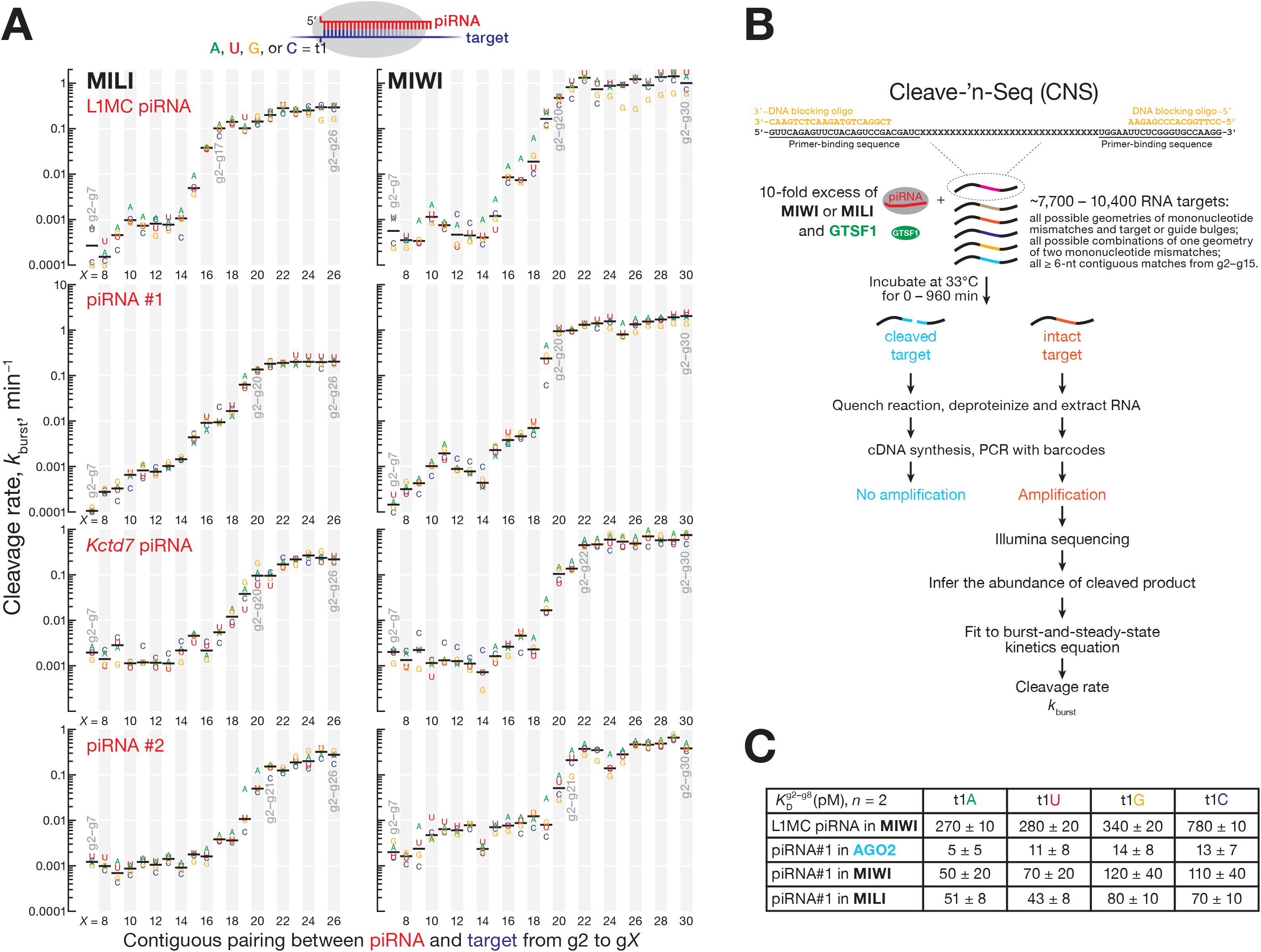
Pairing to piRNA 3’ End is Dispensable for PIWI Slicing, Related to Fig. 2. (A) MILI and MIWI pre-steady-state cleavage rates for targets of piRNAs contiguously paired from nucleotide g2. Data are for targets with all possible identities of nucleotide t1. (B) Overview of Cleave-’n-Seq. (C) AGO2, MILI, and MIWI binding affinities (*K*_D_) for a g2–g8 match with different t1 nucleotide identities. Mean and the range of the data from two independent trials are shown. AGO2 data are from Ref (*39*).

**Fig. S3.**
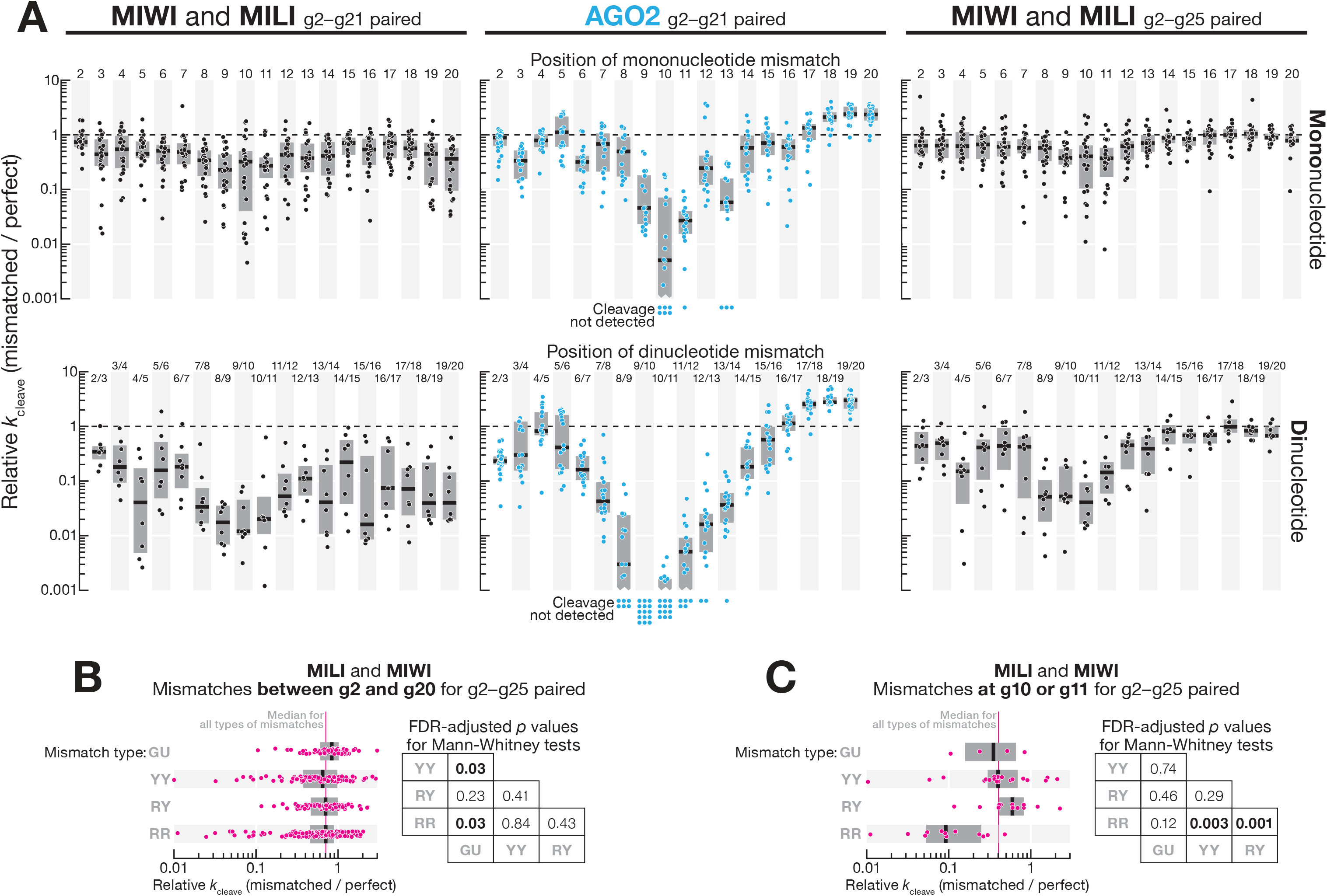
PIWI Slicing Tolerates Mononucleotide and Dinucleotide Mismatches At Any Position, Related to Fig. 2. (A) Change in AGO2, MILI, and MIWI pre-steady-state cleavage rate for one or two consecutive mismatches between g2–g20. Box plots show IQR and median. PIWI data are for all possible mononucleotide mismatch geometries and for one dinucleotide geometry for eight piRISCs (four piRNAs in fig. S1B in MILI and MIWI). AGO2 data are for all possible mononucleotide and dinucleotide mismatch geometries for two RISCs from Ref (*46*). (B) Change in MILI and MIWI pre-steady-state cleavage rate for mismatches between g2–g20. Data are binned by mismatch geometry. Data are for all possible mononucleotide mismatch geometries at all 19 positions between g2–g20 for eight piRISCs (four piRNAs in fig. S1B in MILI and MIWI). Box plots show IQR and median. Kruskal-Wallis test (one-way ANOVA on ranks) *p*-value = 0.025. FDR (Benjamini-Hochberg) corrected *p* values for *post hoc* pairwise Mann-Whitney tests are shown. (C) Change in MILI and MIWI pre-steady-state cleavage rate for mismatches at g10 or g11. Data are binned by mismatch geometry. Data are for all possible mononucleotide mismatch geometries at g10 and g11 for eight piRISCs (four piRNAs in fig. S1B in MILI and MIWI). Box plots show IQR and the median. Kruskal-Wallis test (one-way ANOVA on ranks) *p*-value = 0.001. FDR (Benjamini-Hochberg) corrected *p* values for *post hoc* pairwise Mann-Whitney tests are shown.

**Fig. S4.**
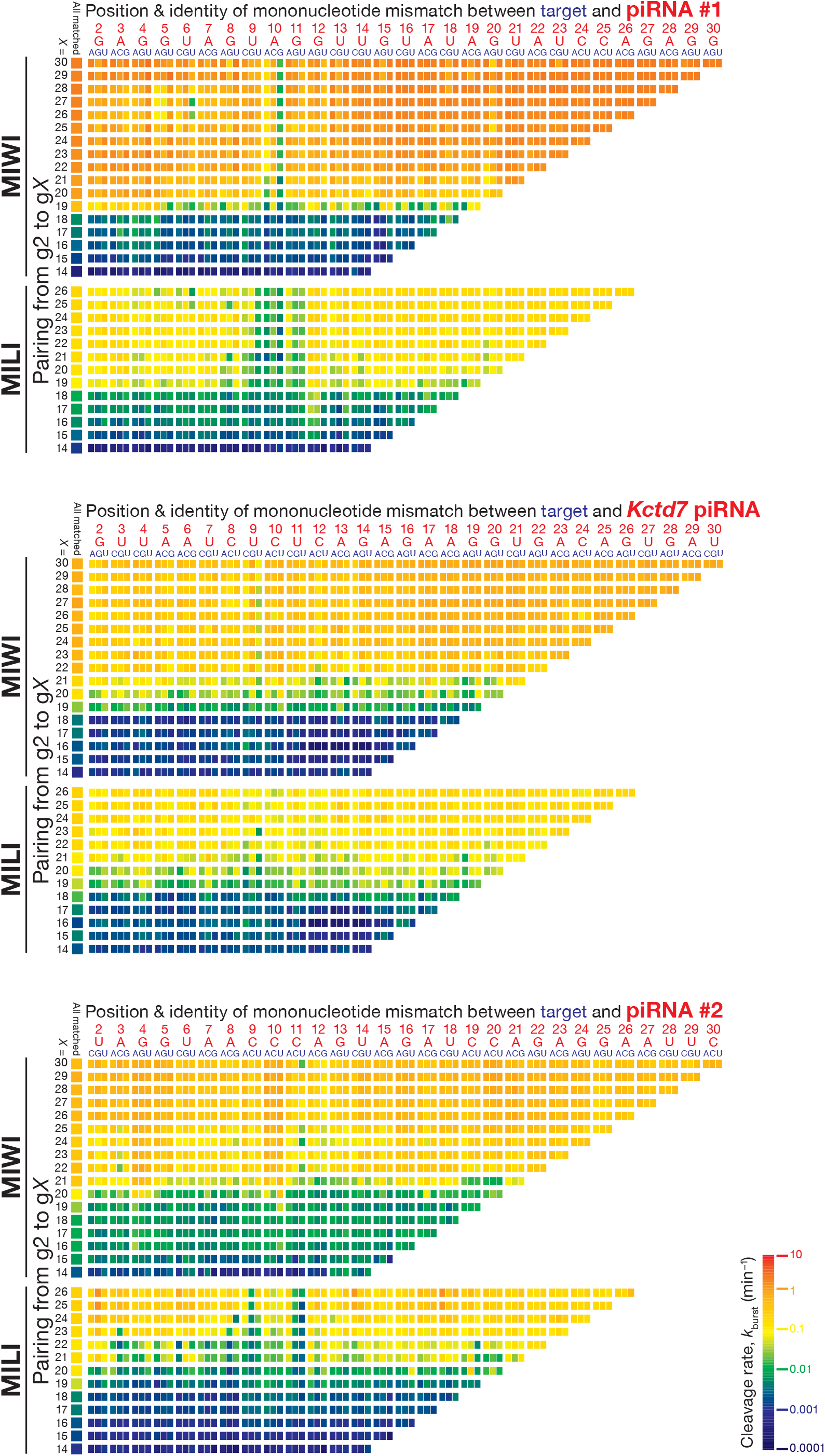
PIWI Slicing Tolerates One Mismatch at Any Position, Related to Fig. 2. MILI and MIWI pre-steady-state cleavage rates for piRNA #1, *Kctd7* piRNA, or piRNA #2 targets containing a single unpaired nucleotide.

**Fig. S5.**
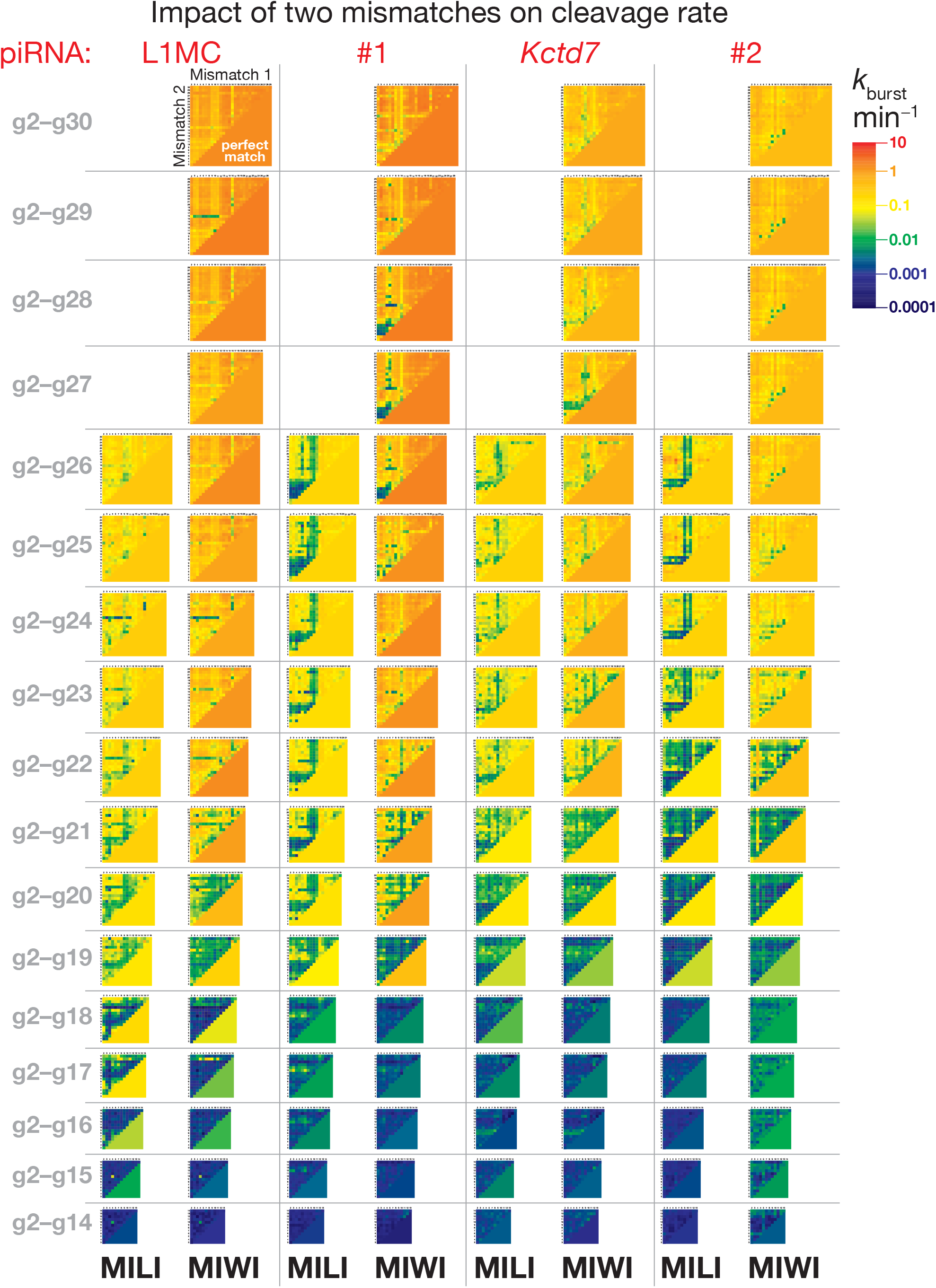
PIWI Slicing Tolerates Two Mismatches at Any Position, Related to Fig. 2. MILI and MIWI pre-steady-state cleavage rates for L1MC piRNA, piRNA #1, *Kctd7* piRNA, or piRNA #2 targets containing two unpaired nucleotides.

**Fig. S6.**
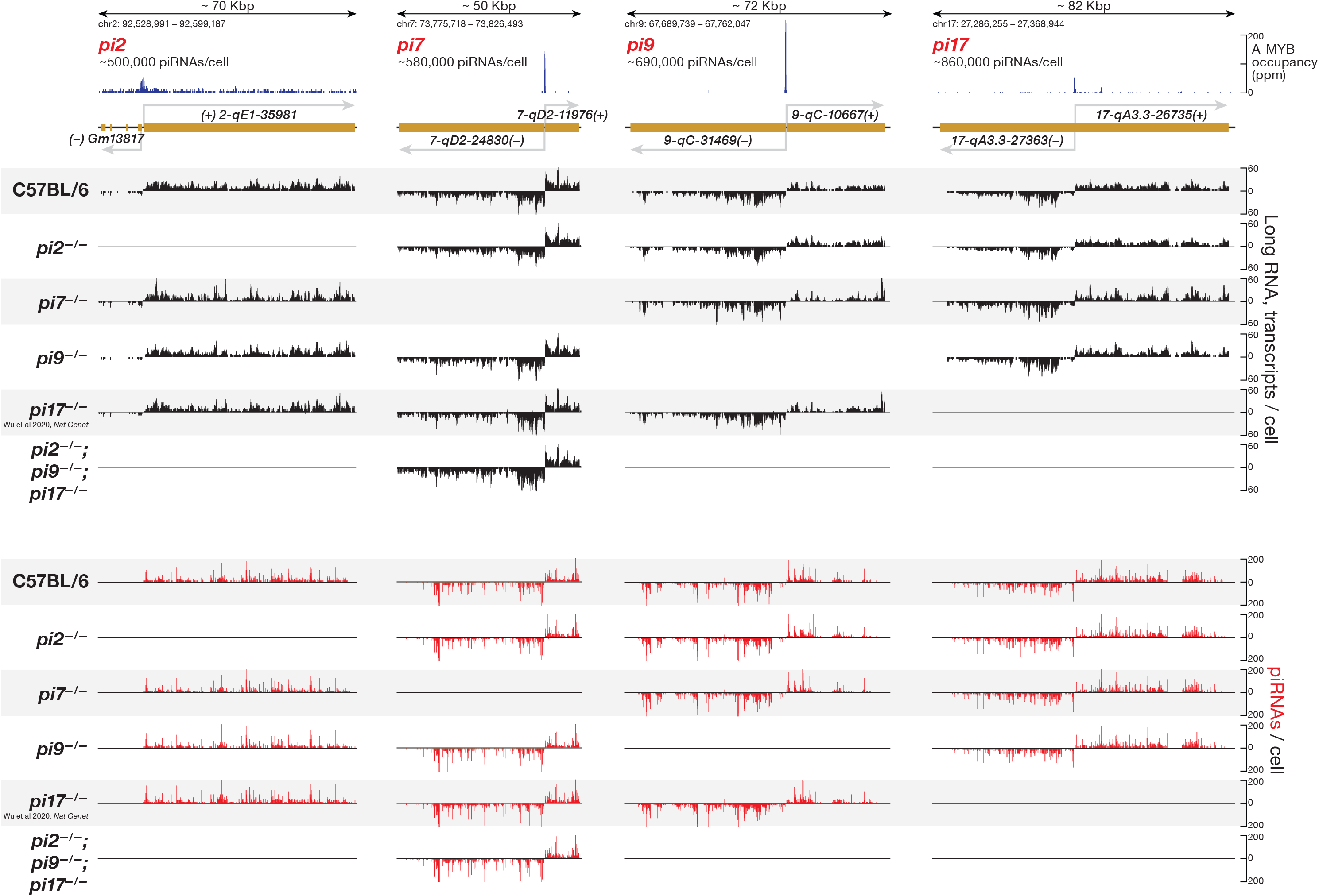
*pi2^−/−^, pi7^−/−^, pi9^−/−^*, and *pi17^−/−^* promoter deletions in mice, Related to Fig. 3. Steady-state abundance of piRNA precursor transcripts and mature piRNAs in FACS-purified mouse primary spermatocytes.

**Fig. S7.**
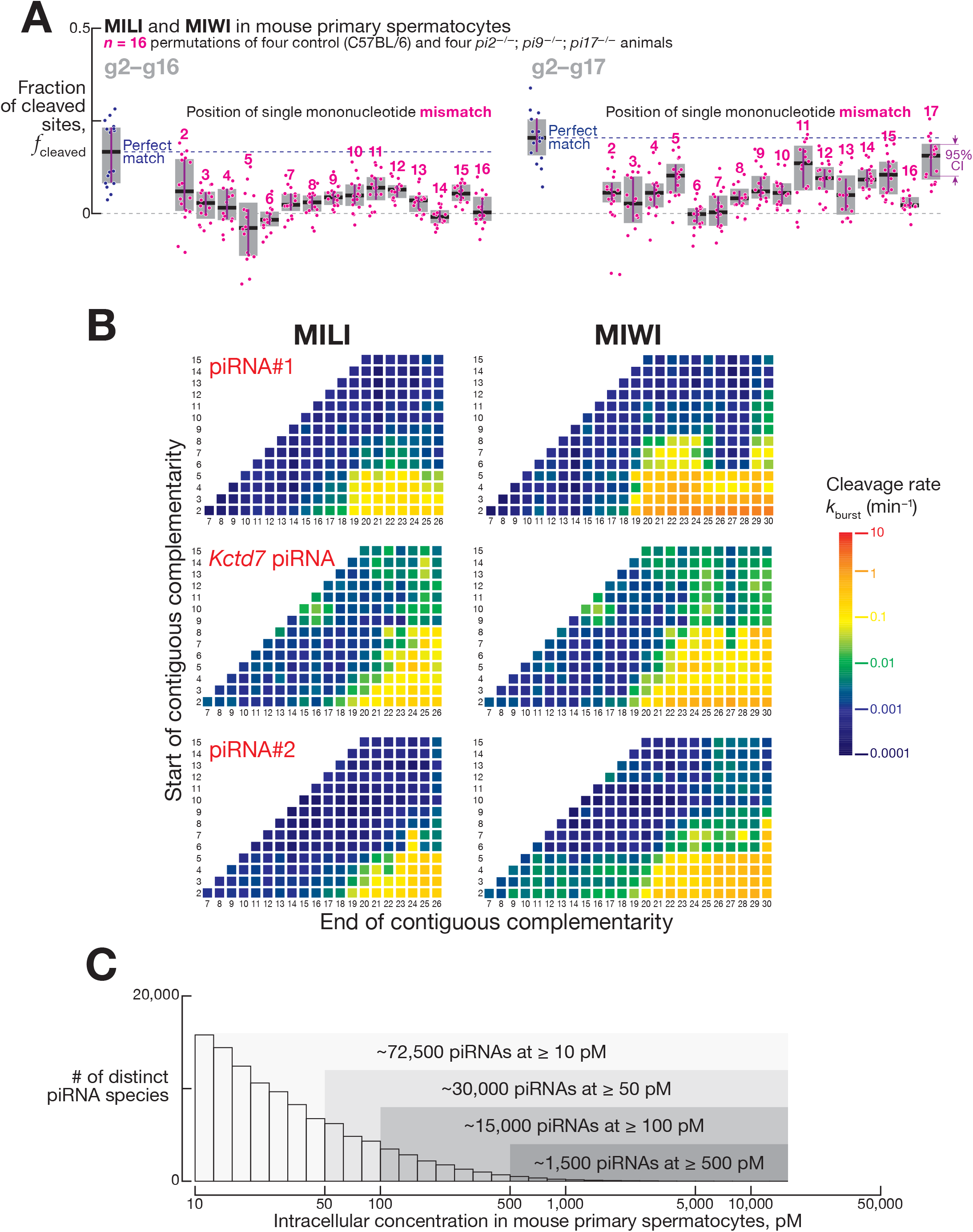
PIWI Slicing Does not Require Pairing to piRNA 5’ End, Related to Fig. 3. (A) Fraction of cleaved targets in FACS-purified mouse primary spermatocytes for pairing containing a single mononucleotide mismatch. Box plots show IQR and median; 95% confidence interval was calculated with 10,000 bootstrapping iterations. (B) MILI and MIWI pre-steady-state cleavage rates in vitro for all possible stretches of ≥ 6-nt contiguous pairing starting from nucleotides g2–g15 of piRNA #1, *Kctd7* piRNA, and piRNA #2. (C) Intracellular concentration of pachytene piRNAs in mouse primary spermatocytes. Data are the mean of 12 biological samples.

**Fig. S8.**
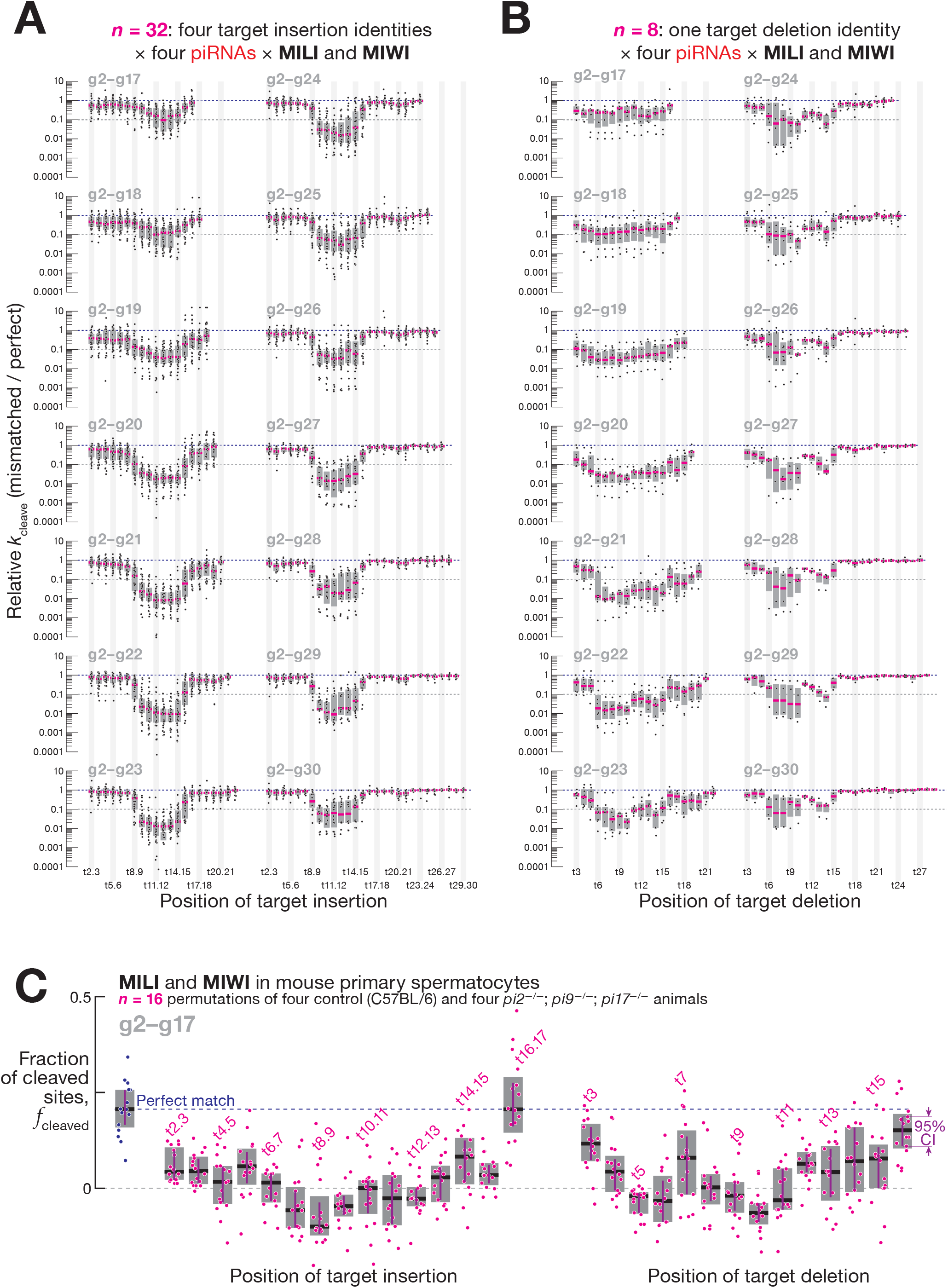
Target Insertions and Deletions in the Center of piRNA:Target Duplex are Detrimental for PIWI Slicing, Related to Fig. 3. (A, B) Change in pre-steady-state cleavage rate for mononucleotide target insertions (A) or target deletions (B). Target insertion data are for 16 or 32 targets (four insertion geometries for four piRNA guides in MILI [for pairing up to g26] or MIWI [for pairing up to g30]). Guide bulge data are for four or eight targets (one deletion geometry for four piRNA guides in MILI [for pairing up to g26] or MIWI [for pairing up to g30]). Box plots show IQR and median. (C) Fraction of cleaved targets in FACS-purified mouse primary spermatocytes for pairing containing a single mononucleotide bulge in target or guide sequence. Box plots show IQR and median; 95% confidence interval was calculated with 10,000 bootstrapping iterations.

**Fig. S9.**
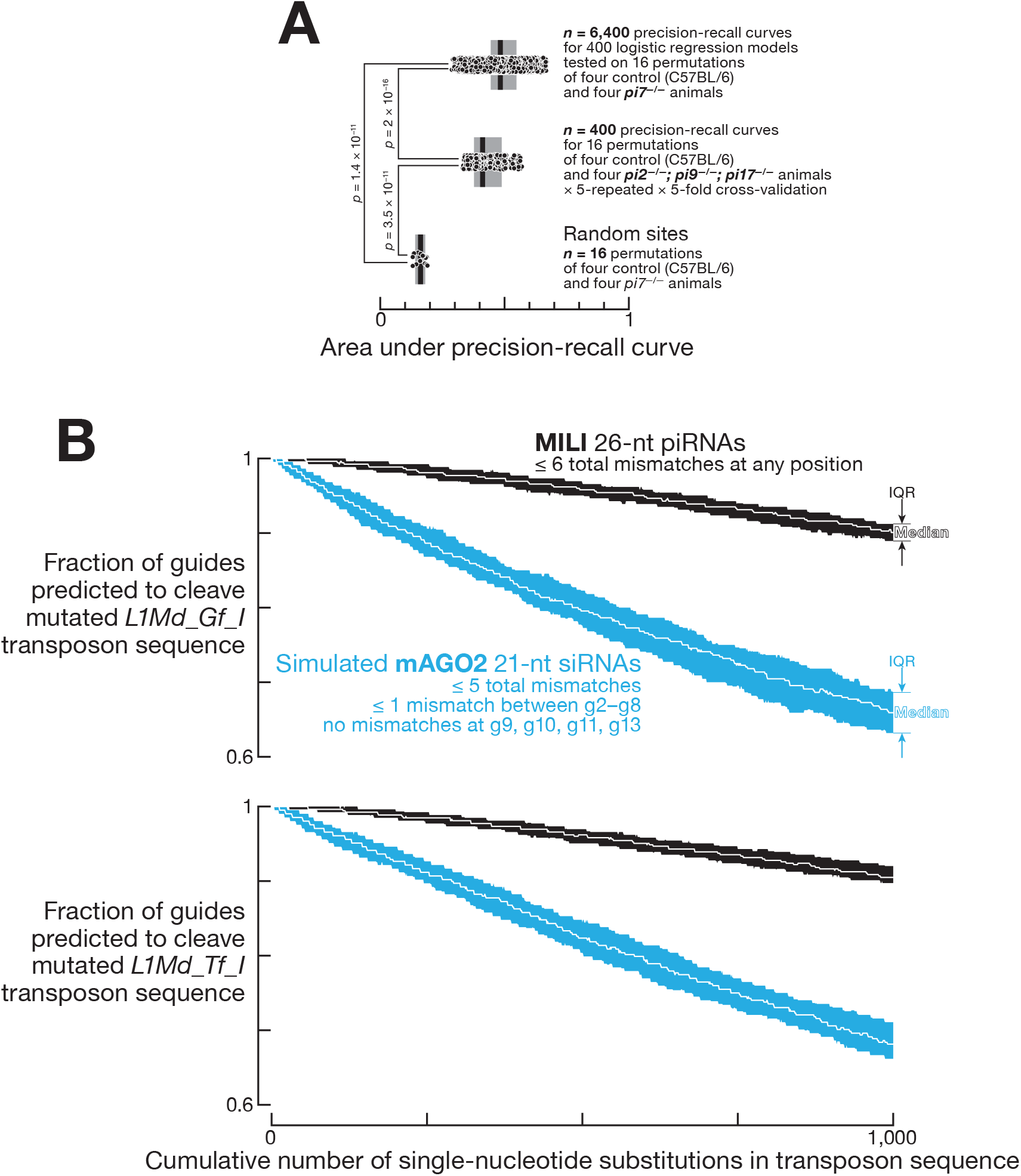
Determinants of Efficient PIWI Slicing, Related to Fig. 4. (A) Area under the precision-recall curves for random control, 400 logistic regression classifier models trained with *pi2^−/−^; pi9^−/−^; pi17^−/−^*; data, and 6,400 tests of the 400 models using *pi7^−/−^* data. Box plots show IQR and median. Kruskal-Wallis test (one-way ANOVA on ranks) *p*-value = 4.5 × 10^-12^. FDR (Benjamini-Hochberg) corrected *p* values for *post hoc* pairwise Mann-Whitney tests are shown. (B) Number of piRNAs and siRNAs predicted to cleave L1Md_Gf and L1Md_Tf transposon sequences. Data are median and IQR from 100 independent simulations.

**Fig. S10.**
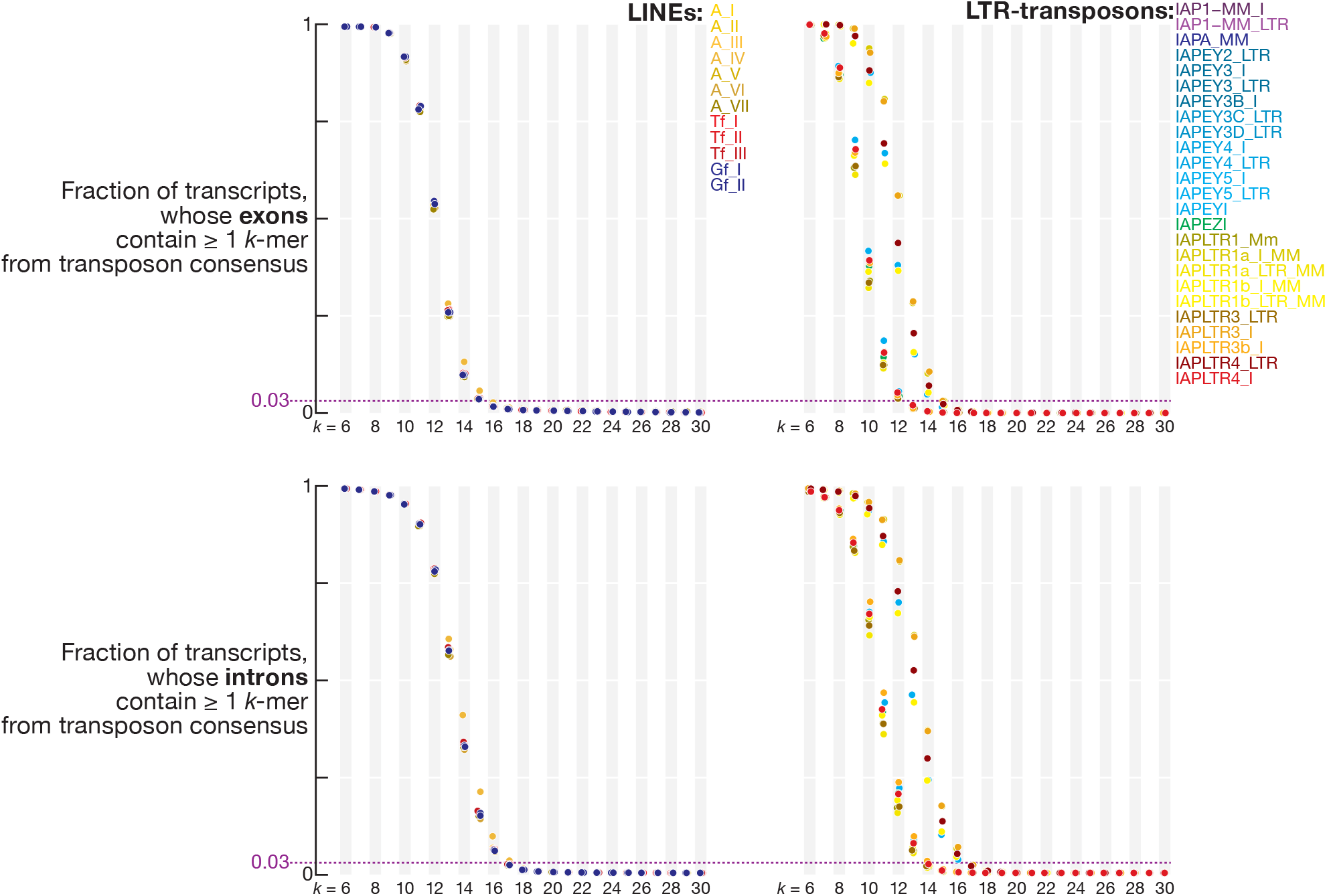
Sequence overlap between transpositionally active families of LINEs and LTR-transposons and exons or introns of mouse mRNAs and lncRNAs.

